# Whole-genome sequencing of *Nicotiana glauca*

**DOI:** 10.1101/211482

**Authors:** Galina Khafizova, Pavel Dobrynin, Dmitrii Polev, Tatyana Matveeva

## Abstract

**Background:** *Nicotiana glauca* (tree tobacco) is a member of the *Solanaceae* family, which includes important crops (potato, tomato, eggplant, pepper) and many medicinal plants. This diploid plant is native to South America and is one of the first *Nicotiana* species with *Agrobacterium* cellular T-DNA (cT-DNA). Its cT-DNA is a partial, inverted repeat, called gT. Tree tobacco belongs to the section *Noctiflorae.* Sequencing of the genomes of *N. tomentosiformis* and *N. otophora* (section *Tomentosae*) and *N. tabacum* (section *Nicotiana*) allowed the detection of previously unknown multiple cT-DNAs, raising the question whether there are other T-DNA insertions in the *N. glauca*. NGS data can help answer this question. Besides, *N. glauca* contains a profile of alkaloids different from *N. tabacum*. The plant is used for medicinal purposes. Comparative analysis of genomic data of phylogenetically distant tobacco species will provide valuable information on the genetic basis for various traits, especially secondary metabolism.

**Findings:** We report a high-depth sequencing and *de novo* assembly of *N. glauca* full genome, which was obtained from 210 Gb Illumina HiSeq data. The final draft genome is 3.2 Gb, with N50 size of 31.1 kbp. T-DNA analysis confirmed the presence of the previously described gT insertion and the absence of other ones.

**Conclusion:** We provide the first comprehensive *de novo* full genome assembly of three tobacco, and a cT-DNA insertion analysis. These genome data could be used in pharmacological and in phylogenetic studies.

## Data Description

### Sample collection

*Nicotiana glauca* seeds were obtained from the State All - Russian scientific research institute of tobacco, makhorka and tobacco products of All - Russian Academy of Agriculture [GNU VNIITTI RAA]. Plant were grown in aseptic conditions on Murashige and Skoog medium [1] and were multiplied by cuttings. Leaf tissue was used for DNA extraction, with a modified version of protocol [2], yielding 30 ng/μl of high molecular weight DNA.

### Library construction

Purified genomic DNA from *Nicotiana glauca* was used to construct both pair-end and mate pair libraries in order to generate a high coverage *de novo* assembly. A pair-end library with an insert size of 350 bp was constructed. For this, the Illumina protocol was used as follows: 1) genomic DNA was fragmented by sonication using an M220 Focused-ultrasonicator; 2) DNA ends were made blunt and an adenine was added to the 3’ ends of the blunt fragments; 3) DNA adaptors [Illumina] with a single “T” were ligated to the DNA fragments above; 4) PCR with 8 cycles was used to enrich DNA fragments with adapters; 5) the library was run on the QIAxcel Advanced System for validation.

To improve resolution of repeats during the assembly stage and scaffolding process, one mate pair library was constructed, with an insert size of 4 kbp. A mate pair library was constructed according to this protocol: 1) genomic DNA was fragmented and tagged by Mate Pair Tagment enzyme according to the Nextera ^®^ Mate Pair Library Prep Reference Guide; 2) DNA fragments were circularized and linear DNA was removed; 3) using Covaris circularized DNA was fragmented again; 4) DNA ends were made blunt and an adenine was added to a 3’ ends of blunt-ended fragments; 5) DNA adaptors [Illumina] with a single “T” were ligated to the DNA fragments above; 6) PCR with 15 cycles was used to enrich those DNA fragments with adapters on both ends; 7) the library was run on a QIAxcel Advanced System and also on an Agilent Technology 2100 Bioanalyzer for validation.

### Read sequencing, quality analysis and filtering

Pair-end and Mate pair libraries were sequenced on four and two lanes yielding 130 Gb and 80 Gb of raw sequence data respectively. Quality of raw reads was analyzed with the FastQC [3] program. Some of the raw reads contained artificial sequences due to adapter contamination, or had low quality base-calling scores. To account for these factors, we filtered and trimmed raw PE reads with Trimgalore [4] [min size 100bp, -e 0.1 -q 30 -O 1 -a AGATCGGAAGAGC] to obtain a clean usable data set. Sequence reads were considered low quality if the quality score was <30 on the phred scale, meaning that the accuracy of those reads was less than 99.9%. An excess of ‘Ns’ (unknown base pairs) in the raw reads was also a signal of low quality. Mate pair raw reads were processed and splitted with Nextclip [5] [with default parameters] and pairs with junction adapter sequence in at least one read and sequence length of 25bp were additionally filtered with Trimgalore [4] [min size 20bp, -e 0.1 -q 28 -O 1 -a CTGTCTCTTATA]. High quality filtered PE and MP reads were used for further genome assembly.

According to the FastQC report, GC content distribution in raw reads is unimodal with a peak around 40%, which suggests that there was no contamination with bacterial or human DNA.

### Genome assembly

The *Nicotiana glauca* was assembled with the MaSuRCA-3.2.2 genome assembler [6]. To estimate insert size and its standard deviation, PE and MP reads were mapped to *Nicotiana tabacum* genome (TN90 strain) [Table 1]. Assembly was done with the following parameters [supplemental file 1]. During assembly, the program performed the following steps: 1) PE and MP reads error correction 2) Contigs assembly from PE reads 3) Contig scaffolding with MP reads 4) Gap closing. Assembly resulted in 385116 scaffolds, with N50 and L50 of 31.1 kbp and 27293 respectively. Genome size estimation based on K-mer analysis was 2 Gb, while the final size of the assembled genome equaled 3.2 Gb.

**Table 1.**
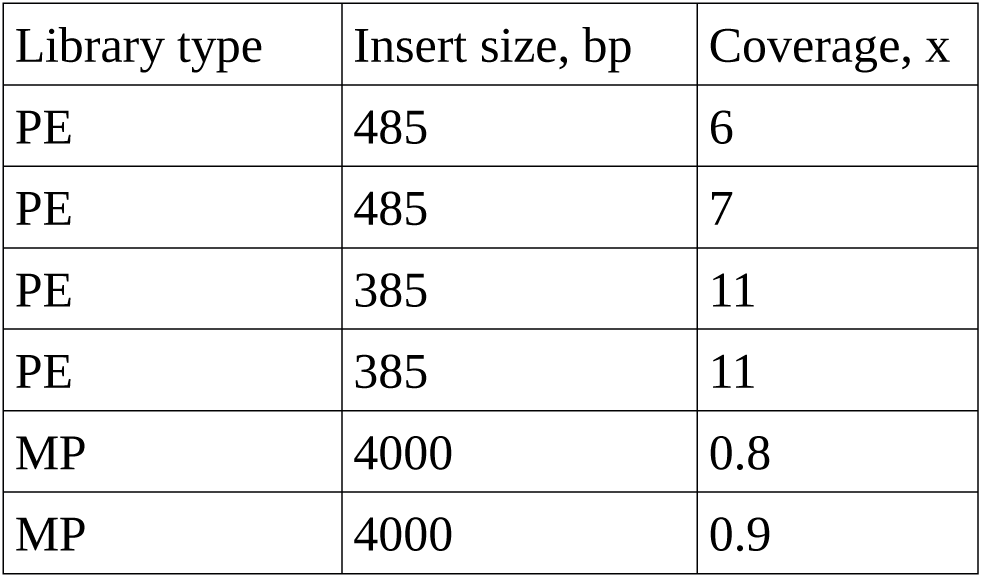
The coverage was measured as the ratio of total size of high-quality filtered reads to the genome size [2 Gb]. Genome size estimation was based on K-mer analysis.

| Library type | Insert size, bp | Coverage, x |
| --- | --- | --- |
| PE | 485 | 6 |
| PE | 485 | 7 |
| PE | 385 | 11 |
| PE | 385 | 11 |
| MP | 4000 | 0.8 |
| MP | 4000 | 0.9 |

### Whole genome alignment of *Nicotiana glauca* and *Nicotiana tabacum*

To identify the location of the *N. glauca* cT-DNA insertion relative to the *N. tabacum* genome, we mapped all *N. glauca* scaffolds to *N. tabacum* scaffolds downloaded from the Sol Genomics Network [7]. To increase accuracy of alignment we masked all known plant repeat classes and their homologs in the *N. glauca* genome. For repeat identification we used the RepeatMasker software [8] and the latest Repbase Update library from 09.27.2017. For whole genome alignment we used the Last software [9]. First we build an index from *N. glauca* scaffolds with the following command: lastdb -P0 -uNEAR -R01 ng-NEAR ng-assembly.fa. After that, substitution and gap frequencies were determined: last-train -P0--revsym --matsym --gapsym -E0.05 -C2 ng-NEAR TN90.fa > ng-TN90.mat. In the final step we build many-to-one alignment of *N. tabacum* to *N. glauca* genome: lastal -m50 -E0.05 -C2-p ng-TN90.mat ng-NEAR TN90.fa | last-split -m1 > ng-TN90.maf.

Comparative analyses of *N. glauca* scaffolds against genome assembly of *N. tabacum* TN90 cultivar strain using the Last software resulted in 3.2 Gbp of aligned sequences with minimum, median and maximum identity of 60%, 88 and 100 % respectively.

### T-DNA analysis

According to our analysis, there are no other T-DNA-like sequences, than those described before [10, 11, 12]. We used the Last software [9] to carry out the alignment of the database [supplemental file 2], to the *N. glauca* genome. The database contained all known T-DNA-like sequences, that were detected as part of cT-DNA [12, 13, 14]. We found sequences homologous to agrobacterial genes *orf13a, orf13, orf14, rolC, rolB* and *mis*. The sequences found form a single region of length 8180 bp and are located on the contig jcf7180010713632. The fragment of T-DNA obtained in the assembly is organized in an imperfect inverted repeat. The similarity of the nucleotide sequences, that we found, and sequence of gT, previously described by Suzuki [12] was 99%, while its similarity to *Agrobacterium* T-DNA is 77-89%. Based on the contig jcf7180010713632 sequence, we designed primers, external to gT insertion. Sequencing of the PCR fragments, amplified from these primers, reaffirm T-DNA/plantDNA junction areas, which together with genome assembly data confirmed the cT-DNA structure, described by Suzuki and co-authors [10, 11, 12]. PCR was carried out using “LONG PCR enzyme Mix” [Thermo scientific] according to the instructions for the kit.

## Conclusion

We generated a *N. glauca* (tree tobacco) genome assembly with T-DNA analysis. Here we confirmed the structure of previously described gT insertion, and demonstrated the absence of new types of cT-DNA in the genome of *N. glauca.* The *de novo* assembled genome of *N. glauca* will provide a valuable resource for pharmacological studies and comparative genomic studies in *Nicotiana* species.

## Abbreviations

T-DNA: transferred DNA
PE: pair-end
MP: mate pair

## Declarations

### Authors’ contributions

TM developed the overall project design. GK, PD and TM wrote the paper. GK collected the *N.glauca* sample and extracted DNA from the sample. GK and DP constructed libraries. DP sequenced the genome of *N. glauca*. PD assembled the *N. glauca* genome and analyzed whole genome alignments. GK, PD and TM performed cT-DNA analysis. All authors read and approved the final manuscript.

## Acknowledgements

The authors thank Professor Otten (Institut de Biologie Moléculaire des Plantes, Strasbourg) and Professor Lutova (Saint Petersburg University, St.Petersburg) for useful discussion and critical reading of the manuscript.

